# Whole-body Microbiota of Newborn Calves and Their Response to Prenatal Vitamin and Mineral Supplementation

**DOI:** 10.1101/2023.03.21.533572

**Authors:** Sarah M. Luecke, Devin B. Holman, Kaycie N. Schmidt, Katherine E. Gzyl, Jennifer L. Hurlbert, Ana Clara B. Menezes, Kerri A. Bochantin, James D. Kirsch, Friederike Baumgaertner, Kevin K. Sedivec, Kendall C. Swanson, Carl R. Dahlen, Samat Amat

**Affiliations:** Department of Microbiological Sciences, North Dakota State University, Fargo, ND, 58108, United States; Lacombe Research and Development Centre, Agriculture and Agri-Food Canada, Lacombe, AB, Canada; Department of Animal Sciences, and Center for Nutrition and Pregnancy, North Dakota State University, Fargo, ND, 58102, United States; Central Grasslands Research Extension Center, North Dakota State University, Streeter, ND, 58483, United States

**Keywords:** newborn, whole-body microbiota, maternal, vitamin and mineral supplementation, bovine

## Abstract

Here, we investigated whether initial microbial colonization at seven different anatomical locations in newborn calves and their blood cytokines are influenced by prenatal vitamin and mineral (VTM) supplementation. Samples were collected from the hoof, liver, lung, nasal cavity, eye, rumen (tissue and fluid), and vagina of beef calves that were born from dams that received diets with or without VTM supplementation throughout gestation (n=7/group). Calves were separated from their dams immediately after birth and fed colostrum and milk replacer until euthanasia at 30 h post-initial colostrum feeding. The microbiota of all samples was assessed using 16S rRNA gene sequencing and qPCR. 15 cytokines and chemokines were quantified in their serum. The hoof, ocular, liver, respiratory, and reproductive sites of newborn calves were colonized by site-specific microbiota that differed from that of the rumen (0.64 ≥ R^2^ ≥ 0.12, *P* ≤ 0.003). Only the ruminal fluid microbiota was differed by on prenatal VTM supplementation (*P<*0.01 Differences (*P*<0.05) were detected in microbial richness (vagina), diversity (ruminal tissue and fluid, eye), composition at the phylum and genus level (ruminal tissue and fluid, and vagina), and total bacterial abundance (ocular and vagina) between VTM and control calves. The cytokine IP-10 was higher (*P*=0.02) in VTM calves. Overall, our results suggest that despite immediate separation from the dam upon birth, whole-body of 32-h old calves are colonized by relatively rich, diverse and site-specific bacterial communities, and that initial microbial colonization of the rumen, vagina and oculus seem to be influenced by the prenatal VTM supplementation.

**IMPORTANCE:** Increased appreciation of maternal nutrition and microbiome’s involvement in developmental programming and evidence supporting *in utero* microbial colonization highlight that maternal nutrition factors could impact offspring microbial colonization. Here, we investigated whether initial microbial colonization in any of 7 different anatomical sites of newborn calves was influenced by maternal vitamin and mineral (VTM) supplementation. We identified changes in ruminal, vaginal, and ocular microbiota in newborn calves in response to prenatal VTM supplementation. We provided a “holistic” view on the whole-body calf microbiota. Our data was obtained from calves of the same sex and age, and who were immediately separated from dams, and hence provides novel insights on taxonomic composition of initial bacterial microbiota colonization in those anatomical sites examined. Combined, this study provides direction for future work targeting the manipulation of early life microbiome via alteration of maternal nutrition and harnessing early life microbiota for improved cattle health and production.

## INTRODUCTION

The global cattle industry is continuously facing complex and multifaceted challenges. These challenges include: 1) increasing antimicrobial resistance, 2) growing concerns over the ecological footprint of meat and milk production, and most importantly, 3) meeting the elevated consumer demand for beef, which is projected to only increase with the global population. Therefore, it is essential that the cattle industry continues to work toward improving feed utilization, reproductive efficiency, and host resilience against infectious diseases, while also reducing the environmental impact of beef and dairy production. Host genetic selection has been a primary target for improving animal health and productivity over the last several decades resulting in tremendous progress in both dairy and beef cattle production systems. However, only recently has the microbiota that colonize various mucosal surfaces of cattle become a target for manipulation/engineering to improve animal health and production (1, 2).

The microbial communities residing within the respiratory (3), gastrointestinal (4), and reproductive tracts (5, 6), are vital to cattle health and productivity as they mediate nutrient metabolism, modulate the immune system, and provide colonization resistance against pathogens when they maintain their symbiotic relationship with the host. In addition to the microbiota present in these three anatomical sites, recent developments highlight that commensal microbiota associated with the oculus (7), hoof (8), and liver (9–12) may mediate bovine ocular, hoof, and liver health. Thus, these microbiota present novel and unique opportunities for the development of microbiome-based strategies to improve feed and reproductive efficiency while also enhancing host resilience against infectious diseases.

However, microbiome manipulation does not lack challenges. For instance, the ruminal (13) and respiratory microbiome (14) in mature animals is resilient and can revert to their original state following the cessation of an intervention. This is advantageous in maintaining a homeostatic microbiome, but it is disadvantageous when introducing new microbes to a mature animal in an attempt to alter the microbiome in a beneficial way. To overcome this challenge, early life microbial programming in young ruminants has been recommended and has shown some efficacy (15–17). Early life microbial programming is based on the current dogma that microbial colonization of the gut starts only during and after birth, and the developing ruminal microbiota within the first three weeks of life is more receptive to manipulation (15). However, emerging evidence from humans (18, 19), bovine fetal fluids (20, 21), the fetal intestine (20–22), as well as the human fetal lung (23), suggests that microbial colonization of calves may begin *in utero.* This, coupled with rodent studies which demonstrate that fetal metabolic and nervous system development is impacted by the maternal microbiota during pregnancy (24, 25), highlight the potential and extended role of the maternal microbiome in calf microbiome development.

While the role of maternal nutrition in programming of offspring metabolic, immune and nervous system development has been well documented in humans and food-producing animals including cattle (26, 27), the potential involvement of the maternal microbiome in the Developmental Origins of Health and Disease (DOHaD) has recently begun to receive greater appreciation (28–31). It has been postulated that maternal dietary alterations may directly or indirectly (via alteration of the maternal gut microbiota) influence fetal development and feto-maternal microbial crosstalk, and that these effects may be transmitted to progeny, thereby altering the offspring microbiota (30, 31).

Vitamin and mineral (VTM) supplementation during pregnancy is an active area of research to identify the link between maternal nutrition and developmental programming of offspring in both humans (32, 33) and cattle (34–36). Vitamin and mineral status play a critical role in the developing fetus, especially during sensitive times of development such as DNA synthesis, tissue growth, neurodevelopment, and organogenesis (32, 34, 37). In the newborn, vitamins and minerals are key players in many metabolic and physiologic processes such as immune function, bone formation, gene expression, and protection from cellular oxidative stress (34, 38). To date, the impact of prenatal VTM supplementation on offspring microbiome development remains underexplored. Given that certain vitamins (e.g. folate, riboflavin, and vitamin K) are synthesized or required by microorganisms (39) (38), and that the gut microbiome influences the bioaccessibility and bioavailability of minerals to the host (40), it is hypothesized that prenatal VTM supplementation may influence early life microbial colonization and offspring microbiome development in cattle.

In the present study, we evaluated whether the bacterial communities colonizing seven different anatomical sites in newborn calf including hoof, liver, lung, nasal cavity, ocular surface, rumen, and vagina are influenced by maternal VTM supplementation throughout the entire gestation. These anatomic sites were selected considering the importance of the microbiome in each of these sites to bovine health and productivity. We also compared the bacterial communities from these different anatomical locations to provide a holistic view of the whole-body biogeography of the bacterial microbiota in the newborn calf. Finally, blood cytokine profiles were compared between VTM and control calves.

## MATERIALS AND METHODS

All experimental procedures were approved by the North Dakota State University (Fargo, ND) Institutional Animal Care and Use Committee (protocol ID: A21047).

### Experimental design and animal husbandry

A schematic overview of the experimental design and sampling regimen is presented in Fig. 1A. Fourteen crossbred Angus-based heifers (initial BW 603.4 ± 2.42 kg) were housed together, individually-fed using a Calan headgate system, and randomly assigned to a basal diet composed of 53% grass hay, 37% corn silage, 10% modified corn distillers grains plus solubles (DDGS) containing 10.63% crude protein (CON; n = 7); or the basal diet supplemented with a vitamin and mineral supplement (VTM; n = 7). The VTM supplement was provided by top dressing 113 g of a loose mineral product (Purina Wind & Rain Storm All-Season 7.5 Complete loose mineral; Land O’Lakes, Inc., Arden Hills, MN) on top of a total mixed ration (TMR) at the time of feeding (Details on the list of VTM presented in Table S1). The diet for both CON and VTM groups was formulated to target gains of 0.45 kg/heifer/day. Heifers were weighed biweekly and fed individually once per day using an electronic head-gate facility (American Calan; Northwood, NH) at the NDSU Animal Nutrition and Physiology Center (ANPC; Fargo, ND).

**Figure. 1.**
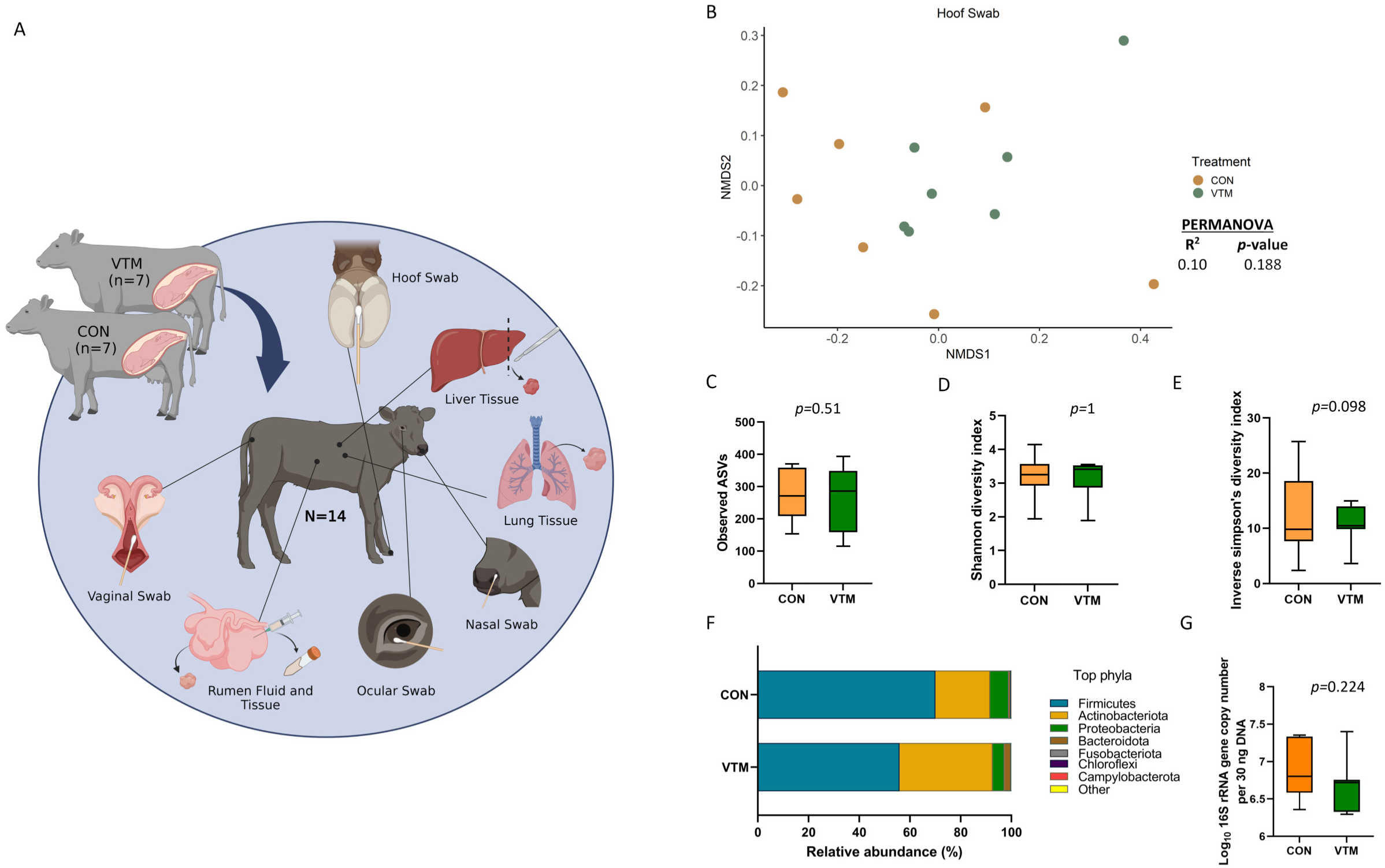
(A) Schematic overview of sampling regimen. Calves were born out of two groups of dams; one group was fed a basal diet plus commercial vitamin and mineral supplement (VTM) and the other was fed only the basal diet (CON) (n = 7/group). Newborn calves from each treatment group were sampled at 30-h after initial colostrum feeding. Samples were collected from each newborn for microbial characterization and consisted of hoof swabs, liver tissue, lung tissue, nasal cavity swabs, ocular swabs, ruminal fluid, ruminal tissue, and vaginal swabs. Created with BioRender.com. (B-G) Hoof microbiota of newborn calves born from dams received either vitamin and mineral (VTM) supplementation or no VTM supplementation received (CON) sampled at 30-h post initial colostrum feeding. (B) Non-metric multidimensional scaling (NMDS) plots of the Bray-Curtis dissimilarities of hoof swab samples; (C) Observed amplicon sequence variants (ASVs) of hoof swab samples; (D) Shannon diversity index of hoof swab samples; (E) Inverse Simpson diversity index of hoof swab samples; (F) relative abundance (%) of top 7 most relatively abundant bacterial phyla of hoof swab samples; (G) total bacterial abundance estimated by qPCR of hoof swab samples.

All heifers underwent a 7-day Select Synch + CIDR estrous synchronization protocol and were bred at the same time point. Heifers were bred on day 60 after initiation of dietary treatments using artificial insemination (AI) of female sexed semen from a single sire. On day 35 post-AI, transrectal ultrasonography was used to determine pregnancy establishment; and this was repeated on day 65 post-Al to confirm that fetuses were female. Later in gestation, heifers were transported to the NDSU Beef Cattle Research Complex (BCRC; Fargo, ND) where they were group-housed and fed individually using the Insentec BV feeding system (Hokofarm Group, Marknesse, The Netherlands). The basal diet (dry matter basis) was composed of 25% corn silage, 66% alfalfa hay, 4% modified corn DDGS, and 5% corn-based premix at 17.5% crude protein and fed for ad libitum intake. All dams remained healthy throughout the study and did not receive any antibiotic treatments.

### Neonatal sample collection

Heifers calved in group pens and calves were immediately removed from the dam without nursing and moved to individual pens. Within 2 h of birth, calves were fed 1.5 L of commercial colostrum replacer that contained 150 g of IgG (LifeLine Rescue High Level Colostrum Replacer, APC, Ankeny, IA, USA) and was delivered using an esophageal tube.

Calves were transported (3 km) from the BCRC to the ANPC and housed indoors and in clean individual pens. Calves were fed 1.9 L of milk replacer (Duralife 20/20 Optimal Non-Medicated Milk Replacer, Fort Worth, TX, USA) via esophageal feeder at 12 and 24 h after the initial feeding. At 30 h after initial colostrum feeding, calves were euthanatized by captive-bolt stunning followed by exsanguination. All sample collections described below were done immediately after euthanasia. Of note, given the logistic challenges associated with calf euthanasia, necropsy and sampling immediately after birth as well as unpredictable timing of calving, euthanasia was performed 30 h after initial colostrum feeding. To remove the confounding effects of dam colostrum and/or milk and milk composition, and microbial transfer from the dam to calf via nursing and direct contact, all calves were separated from the dam immediately after birth and housed individually and fed with colostrum and milk replacer.

### Hoof, ocular, nasal, and vaginal swab sampling

Samples from the hoof and vagina (Fig. 1A) were collected using sterile cotton tipped swabs (Puritan, Guilford, ME, USA). For the hoof, the skin of the interdigital space was swabbed and immediately placed in a sterile Whirl-Pak bag and placed on dry ice. To collect the vaginal samples, the vulva was wiped with 70% ethanol and a paper towel. A sterile cotton swab was inserted into the cranial vagina, swirled four times, carefully removed, and placed into a Whirl-Pak bag and placed on ice. The ocular swab was collected by briefly wiping the outer eyelid with an alcohol prep pad, followed by swabbing of the cornea and conjunctival tissues using an eSwab collection swab (Copan Diagnostics, Murrieta, CA, USA) which was placed into Amies transport media and placed on ice. The nasal swab was collected using an eSwab as well, by briefly inserting the swab into the nostril and swirling against the nasal cavity. These swabs were also placed into Amies transport media and placed on ice until transport to the lab.

### Ruminal fluid, ruminal tissue, liver, and lung tissue sampling

Immediately after euthanasia, the calf was dissected for sample collection. Ruminal fluid was collected from the undeveloped rumen using a sterile 20 ml Luer-Lok syringe with a sterile 18-gauge needle and was immediately transferred into 50-ml Falcon tubes and placed on dry ice. A small portion of the ruminal tissue (ventral sac) was dissected and placed in a sterile microfuge tube and placed on dry ice. The liver and lung were removed, and tissue samples were obtained using a sterile scalpel that was disinfected with an alcohol prep pad between each sample. Tissue samples were placed into individual sterile petri dishes and rinsed with sterile phosphate-buffered saline (PBS) before being aliquoted into sterile microfuge tubes and 1.0 ml stocks of brain heart infusion (BHI) broth containing 20% glycerol. Samples were then placed in dry ice, transferred to the laboratory, and stored at -80 °C until DNA extraction.

### Genomic DNA extraction

Genomic DNA was extracted from the nasal, vaginal, hoof, and ocular swabs using the Qiagen Dneasy Blood and Tissue Kit (Qiagen Inc., Germantown, MD, USA) according to manufacturer instructions with minor modifications. Briefly, the cotton swab tips were submerged in individual aliquots of enzymatic lysis buffer mixture until thoroughly soaked. The enzymatic lysis buffer (20 mM Tris-HCl [pH 8.0], 2 mM sodium EDTA, and 1.2% Triton X-100), also contained 100 mg/ml lysozyme and 25,000 U/ml mutanolysin. The cotton bud was removed from the swab handle using sterilized tweezers and placed back into the enzymatic lysis buffer tube. The samples were then thoroughly vortexed to ensure that the entire cotton swab was submerged and then incubated for 1 h at 37 °C at an agitation rate of 800 rpm. Following this incubation, 25 µl of proteinase K and 400 µl of Buffer AL (without added ethanol) were added to the samples and vortexed. The samples were then incubated for 30 min at 56°C with agitation at 800 rpm. Approximately 400 mg of 0.1 mm zirconia-silica beads were then added to the samples and placed in a FastPrep-24 Classic bead beater (MP Biomedicals, Irvine, CA, USA) and mechanically lysed at 6.0 m/s for 40 s. Samples were then centrifuged at 13,000 × *g* for 5 min. Up to 400 µl of supernatant was removed and placed in a new sterile tube. Then 400 µl of 100% ethanol was added and vortexed. All subsequent steps were following according to the DNeasy Blood and Tissue Kit protocol. Of note, this blood and tissue kit coupled with the enzymatic lysis and bead beating method was used to effectively lyse cells and maximize the DNA extraction from both Gram-positive and Gram-negative species for bacterial community analysis (41).

DNA extraction of the ruminal fluid was performed using the Qiagen DNeasy PowerLyzer PowerSoil kit (Qiagen Inc., Germantown, MD, USA) following the manufacturer’s instructions with modifications previously described (42). DNA was extracted from tissue samples including ruminal tissue, liver tissue, and lung tissue using the Qiagen DNeasy Blood and Tissue Kit (Qiagen Inc., Germantown, MD, USA) with an added bead beating step. After samples were enzymatically lysed, approximately 300 mg of sterile 0.1 mm zirconia-silica beads were added to the samples in bead beating tubes. 200 µl of buffer AL was then added, and samples were placed in a FastPrep-24 bead beater for 40 s at 6.0 m/s. Samples were then spun at 13,000 × *g* for 5 min. The supernatant was transferred into a sterile microfuge tube and the remaining steps were performed according to manufacturer instructions.

### Controls for potential sampling- and DNA extraction-associated contamination

To address the possibility of contamination at the time of sample collection, negative control swabs were collected. A sterile cotton tipped swab (Puritan) was exposed to the ambient air of the tissue collection room and swirled around to capture airborne contaminants.

Additionally, a sterile cotton tipped swab was used to sample the scalpel blade after it had been cleaned and disinfected with ethanol to collect any contaminants surviving the sterilization process between sample collections. DNA from these negative controls was extracted and subjected to 16S rRNA gene sequencing. To account for any contamination introduced during the DNA extraction process, sterile molecular biology grade water (Corning, Corning, NY, USA) was included as a negative extraction control when a fresh kit was opened. Additionally, a sterile cotton-tipped swab and sterile water were included in an extraction to account for any DNA originating from the cotton tip or wooden handle of the swab. These control samples were included in the qPCR and 16S rRNA gene sequencing.

### 16S rRNA gene sequencing and analysis

Amplification and sequencing of the 16S rRNA gene were performed as described previously (20, 43). A total of 124 samples, including 12 environmental and negative DNA extraction controls for each sample type, were sequenced as described in our previous publication (20). Briefly, the V4 region of the 16S rRNA gene was amplified using the 515F (5’-GTGCCAGCMGCCGCGGTAA-3’) and 806R (5’-GGACTACHVGGGTWTCTAAT-3’) primers. A PCR clamp was also included to block amplification of bovine host DNA (44). The PCR amplification was conducted using the HotStarTaq Plus Master Mix Kit (Qiagen Inc., Germantown, MD, United States). After amplification, PCR products were electrophoresed on a 2% agarose gel to ensure the correct size and band intensity. The 16S rRNA gene amplicons were indexed and pooled in equimolar concentrations and then purified using calibrated AMPure XP beads (Indianapolis, IN, United States). Then, the purified PCR product was used to prepare 16S rRNA gene libraries which were then sequenced on an Illumina MiSeq instrument (Illumina Inc., San Diego, CA, United States) using the MiSeq reagent kit v3 (2 × 300 bp) as per manufacturer’s instructions.

Primer sequences were removed from the 16S rRNA gene sequences using Cutadapt v. 4.1 (45). Reads with no primer sequence, shorter than 215 bp, or longer than 285 bp were discarded. DADA2 v. 1.24.0 (46) in R. 4.2.1 was then used to truncate the forward and reverse reads at 200 bp, merge the paired-end reads (minimum overlap 75 bp), remove chimeric sequences, and assign taxonomy to each merged sequence, referred to as amplicon sequence variants (ASVs), using the naïve Bayesian RDP classifier (47) and the SILVA SSU database release 138.1 (48). ASVs that were classified as chloroplasts, mitochondria or eukaryotic in origin were removed as were ASVs likely to be contaminants. ASVs were considered likely contaminants if they were more abundant in the negative controls on average than within each of the sample types. Certain taxa that have been identified as contaminants in previous studies (e.g., *Burkholderia*-*Caballeronia*-*Paraburkholderia*) (49) and not likely to be host-associated were also removed prior to downstream analyses. In total, 1,088 ASVs were removed as likely contaminants (Supplementary Table S2). The number of ASVs per sample (richness), the Shannon and inverse Simpson diversity indices, and Bray-Curtis dissimilarities were calculated in R using Phyloseq 1.40.0 (50) and vegan 2.6-4 (51). To account for uneven sequence depths, all samples were randomly subsampled to 57,500 reads prior to the calculation of Bray-Curtis dissimilarities and microbial diversity measures. All 16S rRNA gene sequences are available in the NCBI sequence read archive under BioProject PRJNA890307.

### Quantification of bacterial abundance using real time qPCR

Quantitative PCR (qPCR) was used to assess the number of 16S rRNA gene copies in each sample using the method we described previously (6, 20). Briefly, primers 515Fq (5’ - GTGYCAGCMGCCGCGGTAA-3’) and 806Rq (5’ -GGACTACNVGGGTWTCTAAT-3’) were used to amplify the V4 region of the 16S rRNA gene. Each qPCR reaction mixture contained 1 SsoAdvanced Universal SYBR Green Supermix (Bio-Rad Laboratories, Inc., Hercules, CA, United States), 0.4 µM (each) primer, 0.1 µg/µl bovine serum albumin (New England Biolabs, Pickering, ON, Canada), 30 ng of DNA template, and 9.4 µl of molecular biology grade water (Corning, Manassas, VA, United States) in a total volume of 25 µl per reaction. A CFX96 Touch Real-Time PCR Detection system (Bio-Rad Laboratories Ltd.) with the following conditions were used: an initial denaturation at 95 °C for 3 min, followed by 40 cycles at 95 °C for 25 seconds, 50 °C for 30 seconds, and then 72°C for 45 seconds. Standard curves (10^2^–10^8^ gene copies) were generated using the pDrive cloning vector (Qiagen Inc.) containing the PCR product from the 16S rRNA gene. All qPCRs were performed in duplicate with standards (1 µl) and no-template controls (1 µl of nuclease-free H_2_O), as well as a positive control (DNA from a bovine deep nasopharyngeal swab). A melt curve analysis was performed following qPCR amplification to ensure that only target genes were amplified. The copy number was normalized to 30 ng of input DNA.

### Quantification of serum cytokines

Blood samples were collected from each calf at the time of exsanguination using 10.0 ml serum vacutainer tubes (Becton Dickinson HealthCare, Franklin Lakes, NJ), and the serum was separated by centrifugation at 1,500 x g for 20 min at 4 °C. Serum was then aliquoted into individual microfuge tubes and stored at -20 °C until processing. A 120 µl aliquot of serum was subjected to multiplexed quantification of 15 bovine cytokines and chemokines. The multiplexing analysis was performed using the Luminex™ 200 system (Luminex, Austin, TX, USA) by Eve Technologies Corp. (Calgary, AB, Canada). Fifteen markers were simultaneously measured using the Eve Technologies Bovine Cytokine 15-Plex Discovery Assay (MilliporeSigma, Burlington, Massachusetts, USA) according to the manufacturer’s protocol. The 15-plex assay consisted of IFNγ, IL-1α, IL-1β, IL-4, IL-6, IL-8, IL-10, IL-17A, IL-36RA, IP-10, MCP-1, MIP-1α, MIP-1β, TNFα, and VEGF-A. Assay sensitivities of these markers range from 0.05 – 66.51 pg/ml for the 15-plex. Individual analyte sensitivity values are available in the MilliporeSigma MILLIPLEX MAP protocol.

### Statistical analysis

Microbial community structure differences between prenatal VTM supplementation groups and between different sample types were assessed using Bray-Curtis dissimilarities and PERMANOVA (adonis2 function) with vegan in R. Pairwise comparisons of the Bray-Curtis dissimilarities between sample types and treatment groups were done using the pairwiseAdonis v. 0.4 R package with the Benjamini-Hochberg procedure used to correct P-values for multiple comparisons. The number of observed ASVs, Shannon diversity index and the estimated bacterial abundance by qPCR among the eight different sample types were compared using the generalized linear mixed model estimation procedure (PROC GLIMMIX) LSMEANS statement. (ver. 9.4, SAS Institute Inc., Cary, NC, USA). The Shapiro-Wilk test was used to determine whether a dataset follows a normal distribution. Differentially abundant genera between the two groups of prenatal supplementation groups were identified using MaAsLin 2 v. 1.10.0 (52) in R. Only those genera with a relative abundance of 0.1% or greater within each sample type were included in this analysis. Significance was considered at *P* < 0.05.

## RESULTS

### 16S rRNA gene sequencing summary

After processing and quality filtering, the average number of sequences per sample were as follows: 185,261 ± 10,017 (SEM) (hoof), 89,852 ± 10,847 (liver), 369, 539 ± 42,475 (lung), 254,352 ± 32,171 (nasal), 264, 921 ± 20,423 (ocular), 269, 858 ± 16,906 (ruminal fluid), 400,929 ± 30,425 (ruminal tissue) and 223,692 ± 15, 009 (vaginal sample) (Table S3). From these sequences, a total of 854 genera from all samples were identified and classified into 31 different phyla (30 bacterial and 1 archaeal phyla).

### Effect of prenatal VTM supplementation on the hoof microbiota

Overall, the epithelial surface of neonatal calf hooves was colonized by a relatively diverse and rich microbial community (Fig. 1B-G). Community structure (R^2^ = 0.10; *P* = 0.188) (Fig. 1B), richness (total observed ASVs) (Fig. 1C) or Shannon diversity (Fig. 1D) of the hoof microbiota did not differ between VTM and control calves. However, VTM calves tended (*P* = 0.098) to have reduced inverse Simpson’s diversity compared to control calves whose dam did not receive VTM supplementation.

Microbial composition at both phylum and genus level did not differ (*P* > 0.1) between the two groups of calves (Fig. 1F). Interestingly, genera such as *Mannheimia*, *Moraxella,* and *Fusobacterium* include potential pathogens associated with bovine respiratory disease (BRD), liver abscesses, and infectious bovine keratoconjunctivitis (IBK) were among the top 10 most relatively abundant genera in the hoof microbiota (Table S5). The estimated total bacterial abundance (16S rRNA gene copy number) as assessed via qPCR did not differ between the VTM and control calves (*P* = 0.224) (Fig. 1G).

### Effect of prenatal VTM supplementation on hepatic microbiota

Sequencing results revealed the presence of a bacterial microbiota in the liver tissue of newborn calves (Fig. 2). There were 51 (± 4 SEM) ASVs observed across all liver tissue samples, which were classified into 18 different bacterial phyla with Firmicutes (62.4%), Proteobacteria (20.8%), and Actinobacteriota (11.4%) being the most dominant phyla. *Staphylococcus* (41.6%), *Corynebacterium* (6.1%), *Streptococcus* (4.6%), *Bacillus* (4.2%), and *Acinetobacter* (3.9%) were the most relatively abundant genera in the liver microbiota. Microbial community structure (R^2^ = 0.07; *P* = 0.703) (Fig. 2A), richness (observed ASVs) (*P* = 0.70, Fig. 2B), and diversity (inverse Simpson) (*P* = 0.138) as well as the relative abundance of phyla (Fig. 2E) and genera, and total bacterial abundance (*P* = 0.20, Fig. 2F) were not affected by prenatal VTM supplementation.

**Figure 2.**
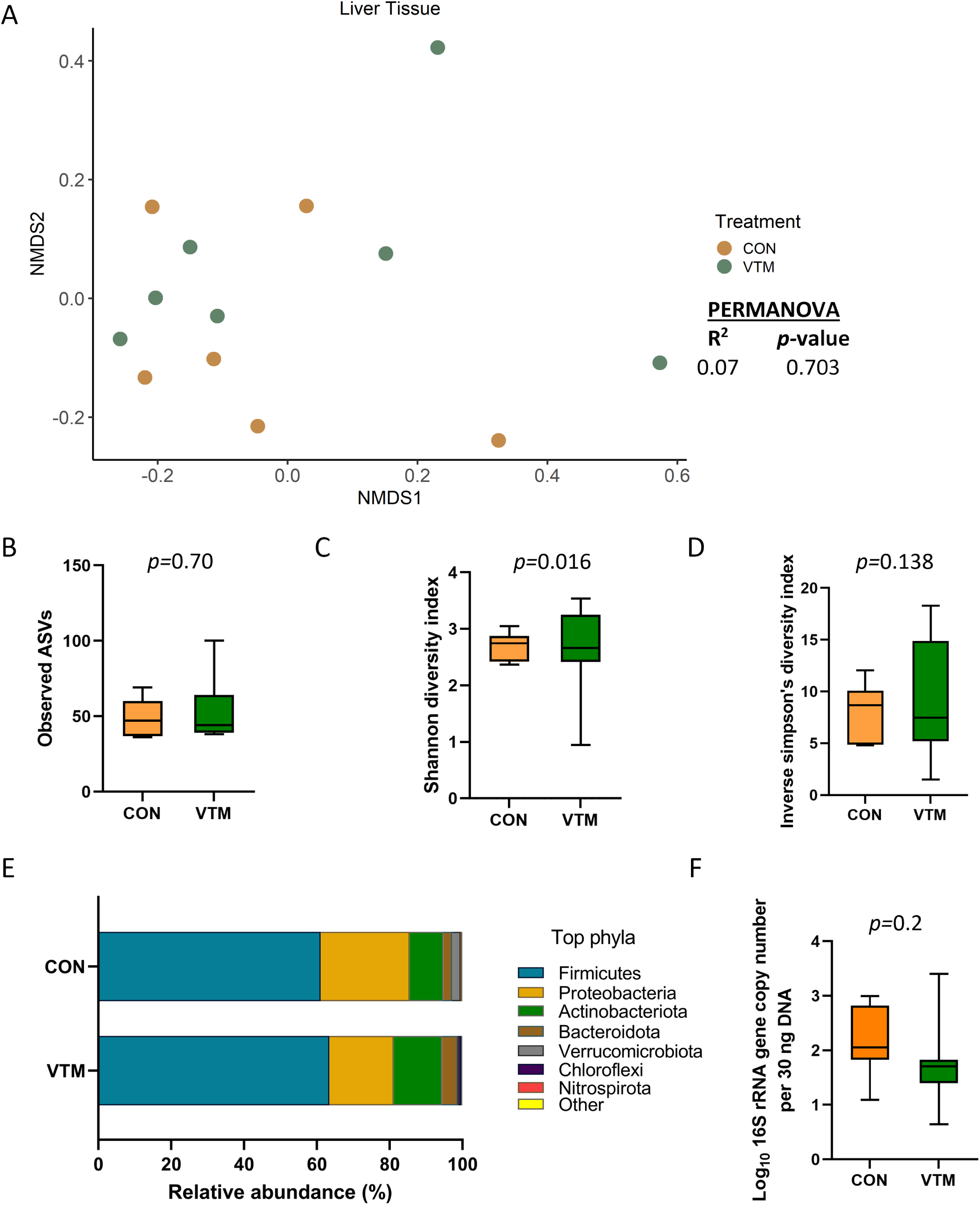
Liver tissue microbiota of newborn calves born from dams received either vitamin and mineral (VTM) supplementation or no VTM supplementation received (CON) sampled at 30-h after initial colostrum feeding (n = 7/group). (A) Non-metric multidimensional scaling (NMDS) plots of the Bray-Curtis dissimilarities; (B) Observed amplicon sequence variants (ASVs); (C) Shannon diversity index; (D) Inverse Simpson diversity index; (E) relative abundance (%) of top 7 most relatively abundant bacterial phyla; (F) total bacterial abundance estimated by qPCR.

### Effect of prenatal VTM supplementation on the respiratory microbiota

A diverse and unique microbial community was detected in the nasal cavity and lung tissue of the newborn calves (Fig 3). The nasal microbiota was not affected by prenatal VTM supplementation as demonstrated by the similar microbial community structure (Fig 3A). Alpha diversity (Fig. 3B-D) measures, phyla composition (Fig. 3E), and estimated total bacterial abundance (Fig. 3F) were the same between the two group calves (*P* > 0.1). Nasal cavity was harbored by respiratory pathogen containing genera *Mannheimia* and *Moraxella* but their relative abundance did not differ between the VTM and control calves (*P* > 0.05). A total of 18 different phyla was detected with Firmicutes (68.7%), Proteobacteria (20%), and Actinobacteriota (7.4%) accounting for more than 95% of the total nasal microbiota. *Streptococcus* (39.5%) and *Staphylococcus* (11.1%) were the most dominant genera.

**Figure 3.**
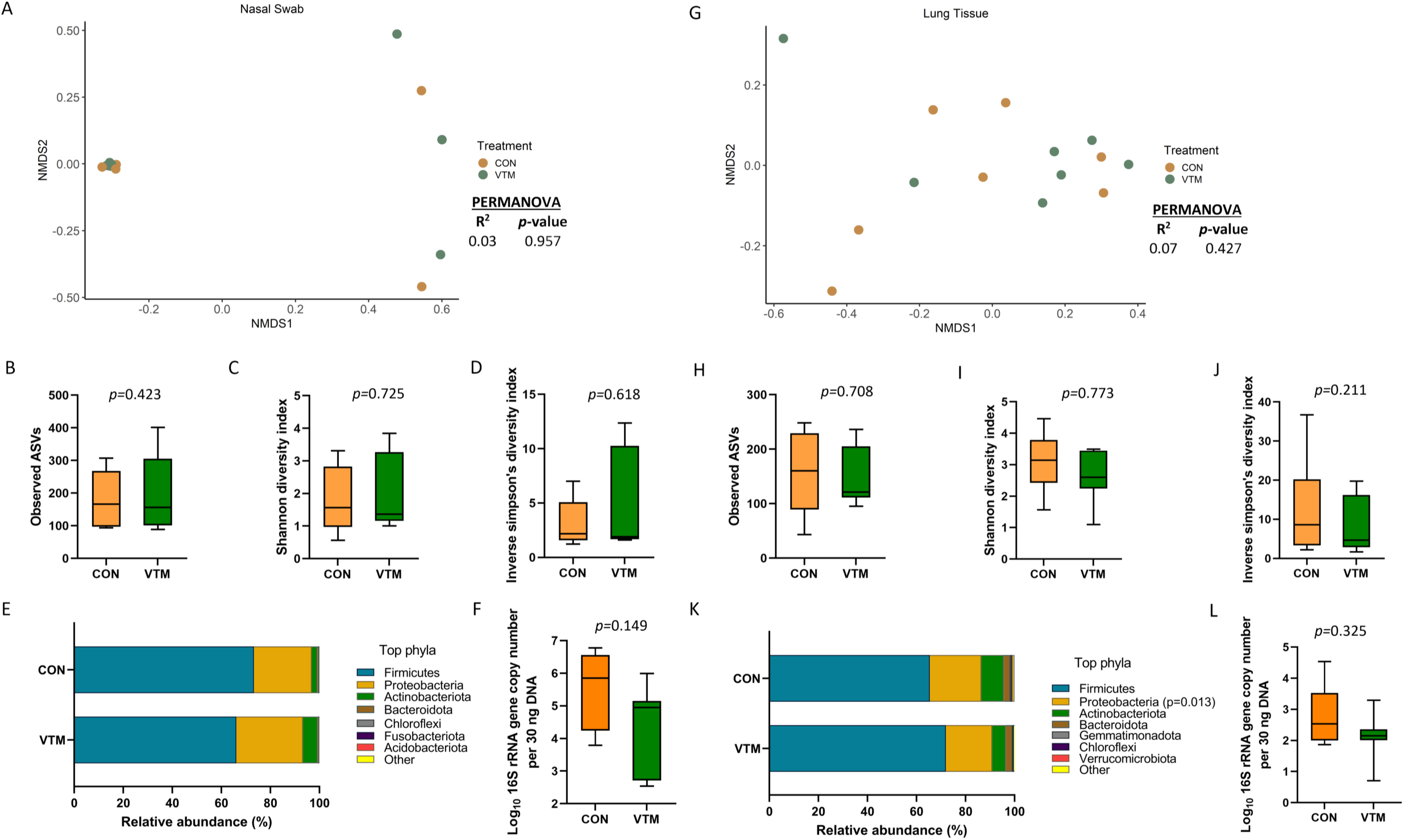
Respiratory microbiota (Nasal [A-F] and lung [G-L] microbiota) of newborn calves born from dams received either vitamin and mineral (VTM) supplementation or no VTM supplementation received (CON) sampled at 30-h after initial colostrum feeding (n = 7/group). (A) Non-metric multidimensional scaling (NMDS) plots of the Bray-Curtis dissimilarities of nasal swab samples; (B) Observed amplicon sequence variants (ASVs) of nasal swab samples; (C) Shannon diversity index of nasal swab samples; (D) Inverse Simpson diversity index of nasal swab samples; (E) relative abundance (%) of top 7 most relatively abundant bacterial phyla of nasal swab samples; (F) total bacterial abundance estimated by qPCR of nasal swab samples. (G) Non-metric multidimensional scaling (NMDS) plots of the Bray-Curtis dissimilarities of lung tissue samples; (H) Observed amplicon sequence variants (ASVs) of lung tissue samples; (I) Shannon diversity index of lung tissue samples; (J) Inverse Simpson diversity index of lung tissue samples; (K) relative abundance (%) of top 7 most relatively abundant bacterial phyla of lung tissue samples; (L) total bacterial abundance estimated by qPCR of lung tissue samples.

Similar to the nasal microbiota, the lung tissue microbial community structure, richness, diversity, and total bacterial abundance (Fig. 3G-J & 3L) did not differ between prenatal treatment groups (*P* ≥ 0.21). However, the relative abundance of the phylum Proteobacteria was lower (*P* = 0.013; Fig. 3K) in VTM calves, no differentially abundant genera between VTM and control calves was detected. The lung tissue of the newborn calves harbored a slightly less rich microbial community (152 ± 17 total ASVs) than the nasal microbiota (185 ± 26).

### Effect of prenatal VTM supplementation on the rumen microbiota

There were only 97 (± 5) total observed ASVs present in the ruminal fluid which were assigned to 7 bacterial phyla. *Firmicutes* (81.1%) and *Proteobacteria* (18.8%) accounted for nearly all (99.9%) of the 16S rRNA gene sequences. A significant effect of prenatal VTM supplementation on the microbial community structure of the ruminal fluid microbiota was detected (*R^2^* = 0.27; *P* = 0.002) (Fig. 4A). Diversity, as measured with the inverse Simpson index, was lower in the ruminal fluid microbiota of VTM calves compared to that of control calves (*P* = 0.021, Fig. 4D). No significant difference was detected in microbial richness (Fig. 4B), Shannon diversity (Fig. 4C), relative abundance of most relatively abundant phyla (Fig. 4E), or the total estimated bacterial abundance (Fig. 4F) (*P* > 0.1). The relative abundance of *Escherichia-Shigella* (*P* = 0.0002) was reduced in VTM calves compared to control calves (Fig. 7).

**Figure 4.**
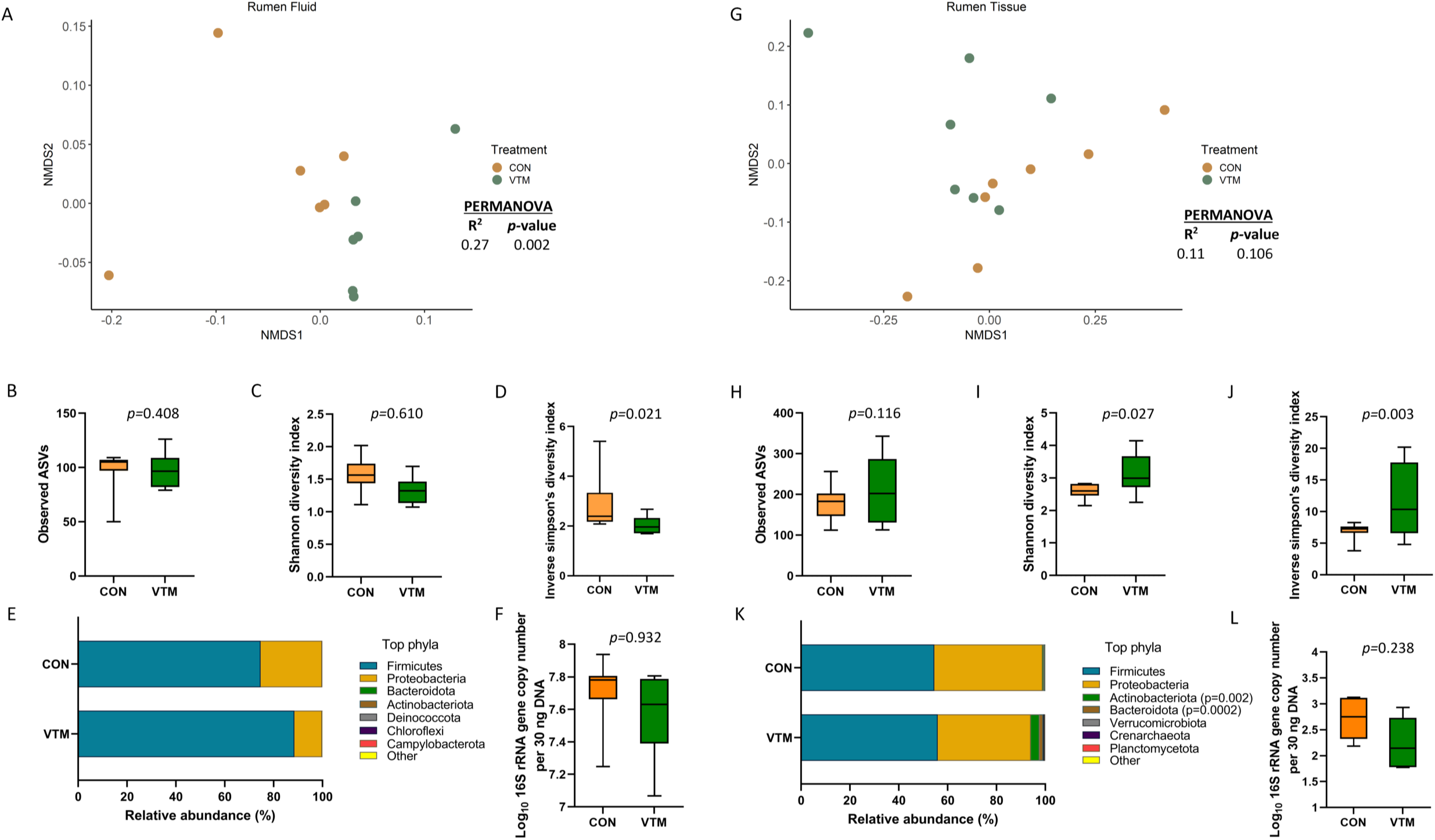
Ruminal microbiota (Ruminal fluid [A-F] and ruminal tissue [G-L] microbiota) of newborn calves born from dams received either vitamin and mineral (VTM) supplementation or no VTM supplementation received (CON) sampled at 30-h after initial colostrum feeding (n = 7/group). (A) Non-metric multidimensional scaling (NMDS) plots of the Bray-Curtis dissimilarities of ruminal fluid samples; (B) Observed amplicon sequence variants (ASVs) of ruminal fluid samples; (C) Shannon diversity index of ruminal fluid samples; (D) Inverse Simpson diversity index of ruminal fluid samples; (E) relative abundance (%) of top 7 most relatively abundant bacterial phyla of ruminal fluid samples; (F) total bacterial abundance estimated by qPCR of ruminal fluid samples. (G) Non-metric multidimensional scaling (NMDS) plots of the Bray-Curtis dissimilarities of ruminal tissue samples; (H) Observed amplicon sequence variants (ASVs) of ruminal tissue samples; (I) Shannon diversity index of ruminal tissue samples; (J) Inverse Simpson diversity index of ruminal tissue samples; (K) relative abundance (%) of top 7 most relatively abundant bacterial phyla of ruminal tissue samples; (L) total bacterial abundance estimated by qPCR of ruminal tissue samples.

The microbiota of the ruminal tissue was more diverse compared to that of the ruminal fluid. The microbial community structure (*R^2^* = 0.11; *P* = 0.106; Fig. 4G), richness (*P* = 0.116; Fig. 4H) and total bacterial abundance (*P* = 0.238; Fig. 4L) in the ruminal tissue did not diverge between the VTM and control calves. However, the ruminal tissue microbiota in the VTM calves was more diverse (*P* ≤ 0.027; Fig. 4I, J) and contained a greater (*P* ≤ 0.002) relative abundance of Actinobacteriota and Bacteroidota phyla compared to the control calves. Among the four genera that were differentially abundant between the VTM and control calves (Fig. 7), *Anaerococcus, Micrococcus* and *Staphylococcus* were significantly enriched (*P* ≤ 0.002) while Escherichia-Shigella (*P* = 0.0009) was reduced in in VTM calves (Fig. 7).

### Effect of prenatal VTM supplementation on the ocular microbiota

The ocular surface was colonized by a rich (total ASVs = 426 ± 36) and diverse microbial community dominated by the phyla Firmicutes (79.7%), Proteobacteria (17.1%), and Actinobacteriota (7.0%). Calves born from VTM supplemented dams had greater microbial diversity (inverse Simpson index; *P* = 0.045; Fig. 5D) yet lower bacterial abundance (*P* < 0.001; Fig. 5F). The relative abundance of the Bacteroidota and Chloroflexi phyla was higher (Fig. 5E; *P* ≤ 0.042) in the VTM calves, and no differentially abundant genera between the two groups was detected. The total bacterial concentration in the ocular samples was lower in the VTM calves (Fig. 5F).

**Figure 5.**
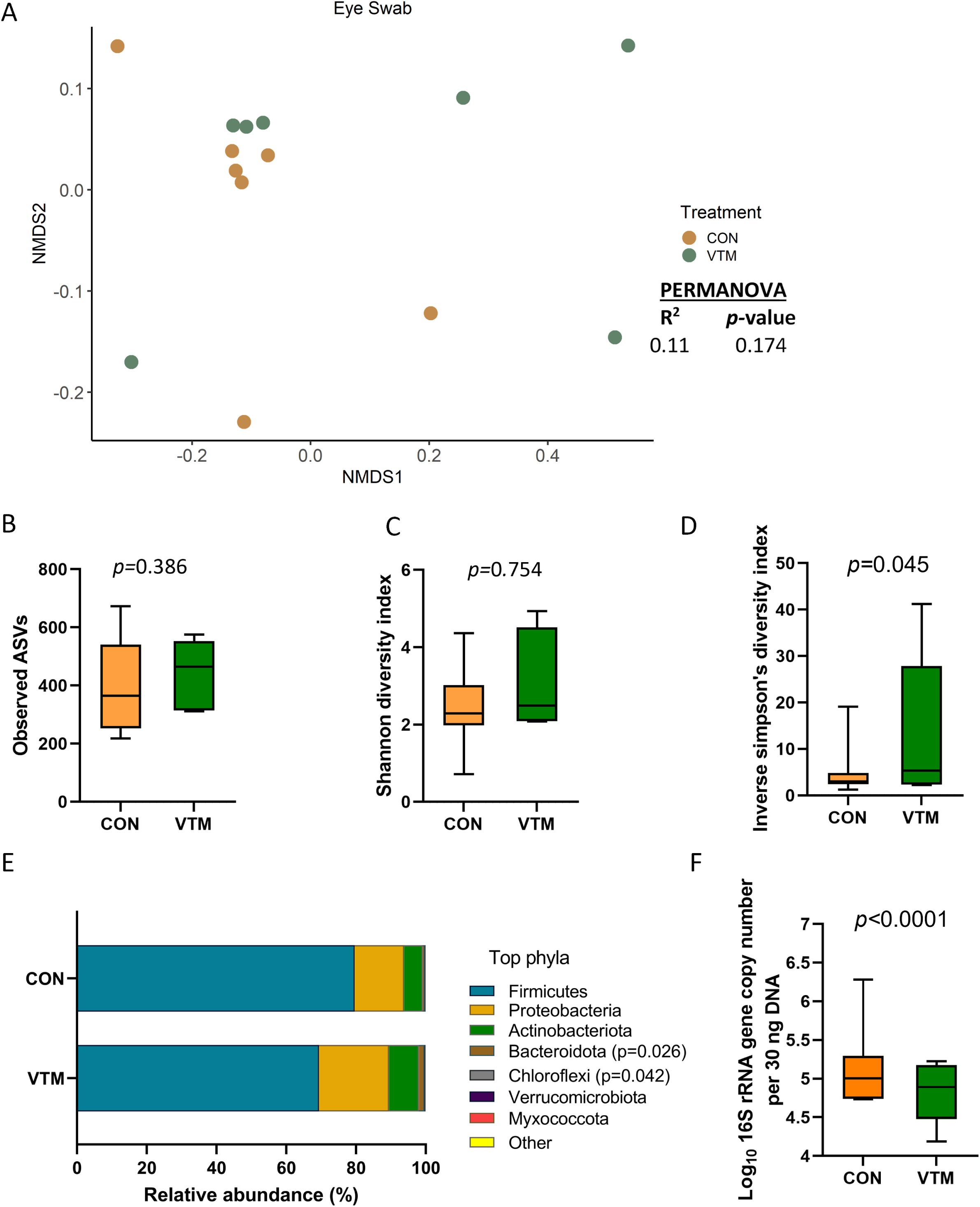
Ocular microbiota of newborn calves born from dams received either vitamin and mineral (VTM) supplementation or no VTM supplementation received (CON) sampled at 30-h after initial colostrum feeding (n = 7/group). (A) Non-metric multidimensional scaling (NMDS) plots of the Bray-Curtis dissimilarities; (B) Observed amplicon sequence variants (ASVs); (C) Shannon diversity index; (D) Inverse Simpson diversity index; (E) relative abundance (%) of top 7 most relatively abundant bacterial phyla; (F) total bacterial abundance estimated by qPCR.

### Effect of prenatal VTM supplementation on the vaginal microbiota

The vagina of the newborn calf was colonized by a rich (total ASVs = 308 ± 46) and diverse (26 different phyla) microbial community. While microbial community structure (*R^2^*= 0.10; *P* = 0.207; Fig. 6A) and diversity (*P* ≥ 0.125) did not differ between groups (Fig. 6C, D), richness (*P* = 0.031) was greater (Fig.6B); while the bacterial concentration (*P* < 0.0001; Fig. 6F) was lower in the vagina of VTM calves compared to control calves. Among the 7 most relatively abundant phyla, the relative abundance of Bacteroidota was greater in the VTM calves (*P* = 0.0008; Fig. 6E). Genera *Corynebacterium*, *Brevibacterium* and *Brachybacterium* were enriched in VTM calves (*P* ≤ 0.005) (Fig. 7).

**Figure 6.**
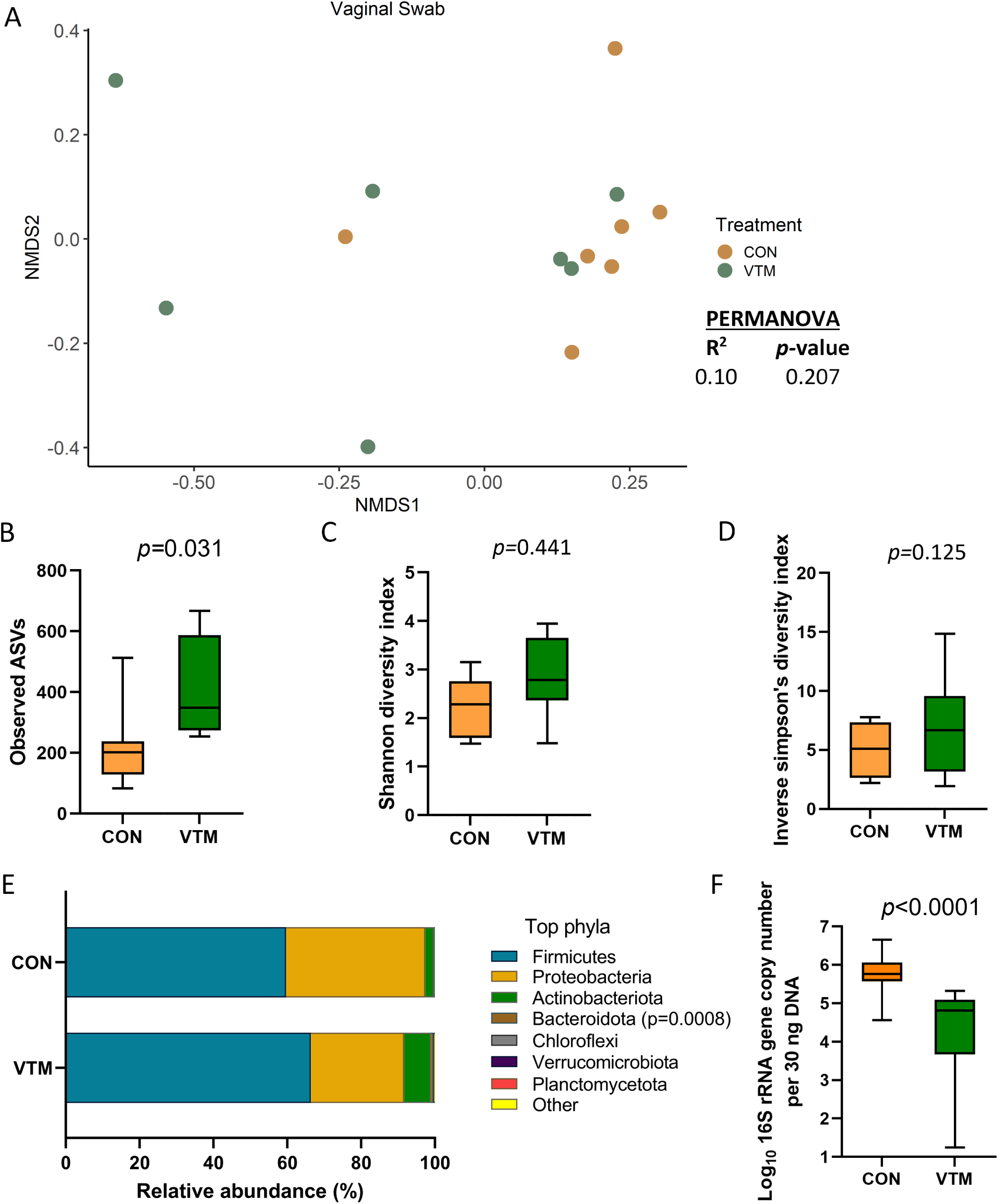
Vaginal microbiota of newborn calves born from dams received either vitamin and mineral (VTM) supplementation or no VTM supplementation received (CON) sampled at 30-h after initial colostrum feeding (n = 7/group). (A) Non-metric multidimensional scaling (NMDS) plots of the Bray-Curtis dissimilarities; (B) Observed amplicon sequence variants (ASVs); (C) Shannon diversity index; (D) Inverse Simpson diversity index; (E) relative abundance (%) of top 7 most relatively abundant bacterial phyla; (F) total bacterial abundance estimated by qPCR.

**Figure 7.**
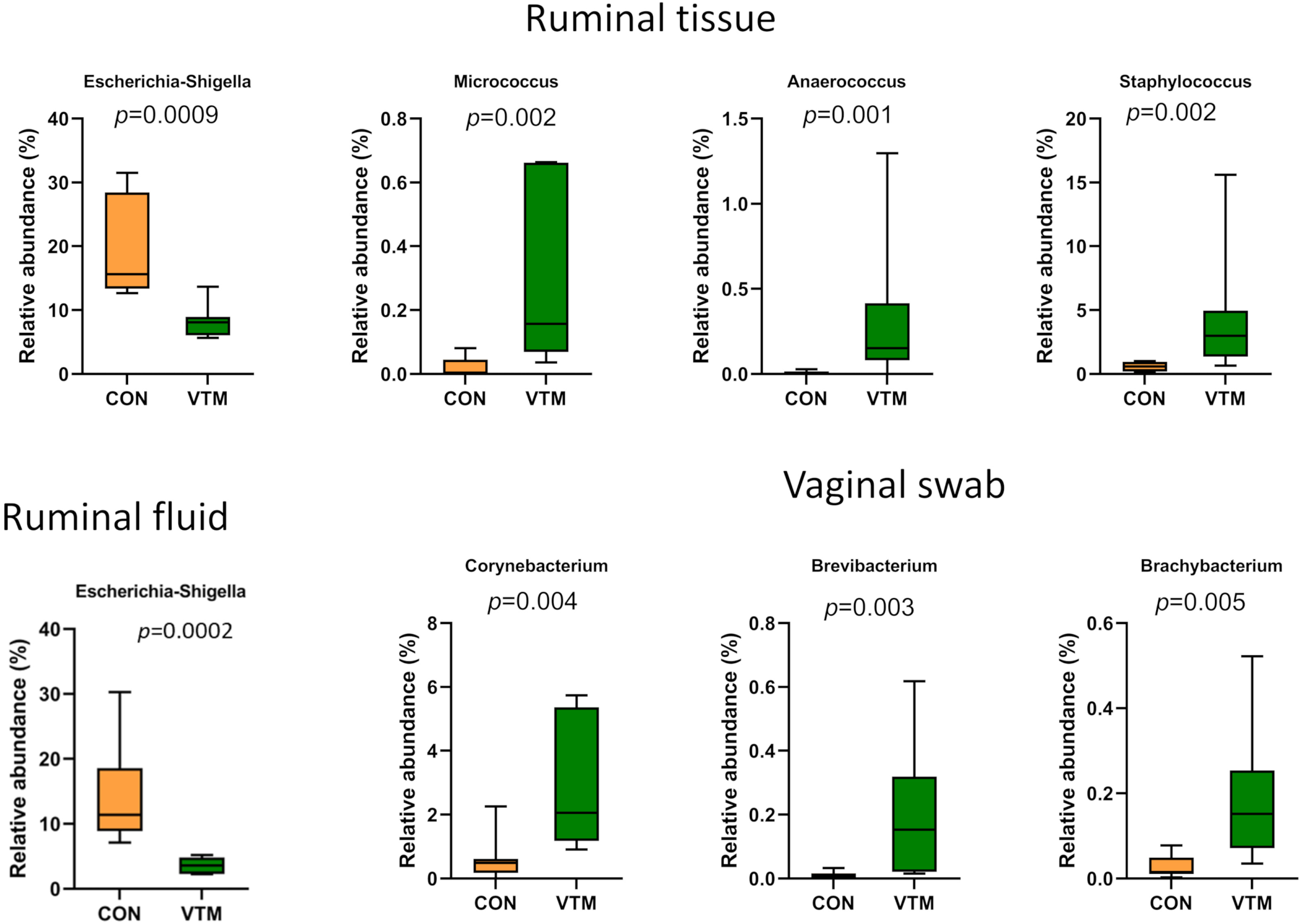
Bacterial genera that were differentially abundant between VTM and control calves (n = 7/group).

### Similarities and differences among the calf microbial communities

The community structures of the hoof, liver, ocular, and vaginal microbiota were different from the rumen-associated microbiota (0.64 ≥ R^2^ ≥ 0.12, *P* ≤ 0.003). The structure of the respiratory (nasal and lung) tract microbiota was similar to that of ocular, vaginal, and ruminal fluid microbiota (*P* > 0.11) (Fig. 8A; Table S4). The ocular, hoof, and vagina had the greatest microbial richness (observed ASVs; *P* < 0.05; Fig.8B) but similar diversity (Shannon; *P* > 0.05; Fig. 8C) compared to other sampling locations. The liver and ruminal fluid had the least microbially rich communities, the latter also being the least diverse (*P*< 0.05). Firmicutes, Actinobacteriota, and Proteobacteria were the dominant phyla across all sample types, but their relative abundance varied by location (Fig. 8E). The ruminal fluid and hoof had the highest bacterial concentration, while the liver, lung, and ruminal tissue had lower bacterial abundance as estimated by qPCR (Fig. 8D).

**Figure 8.**
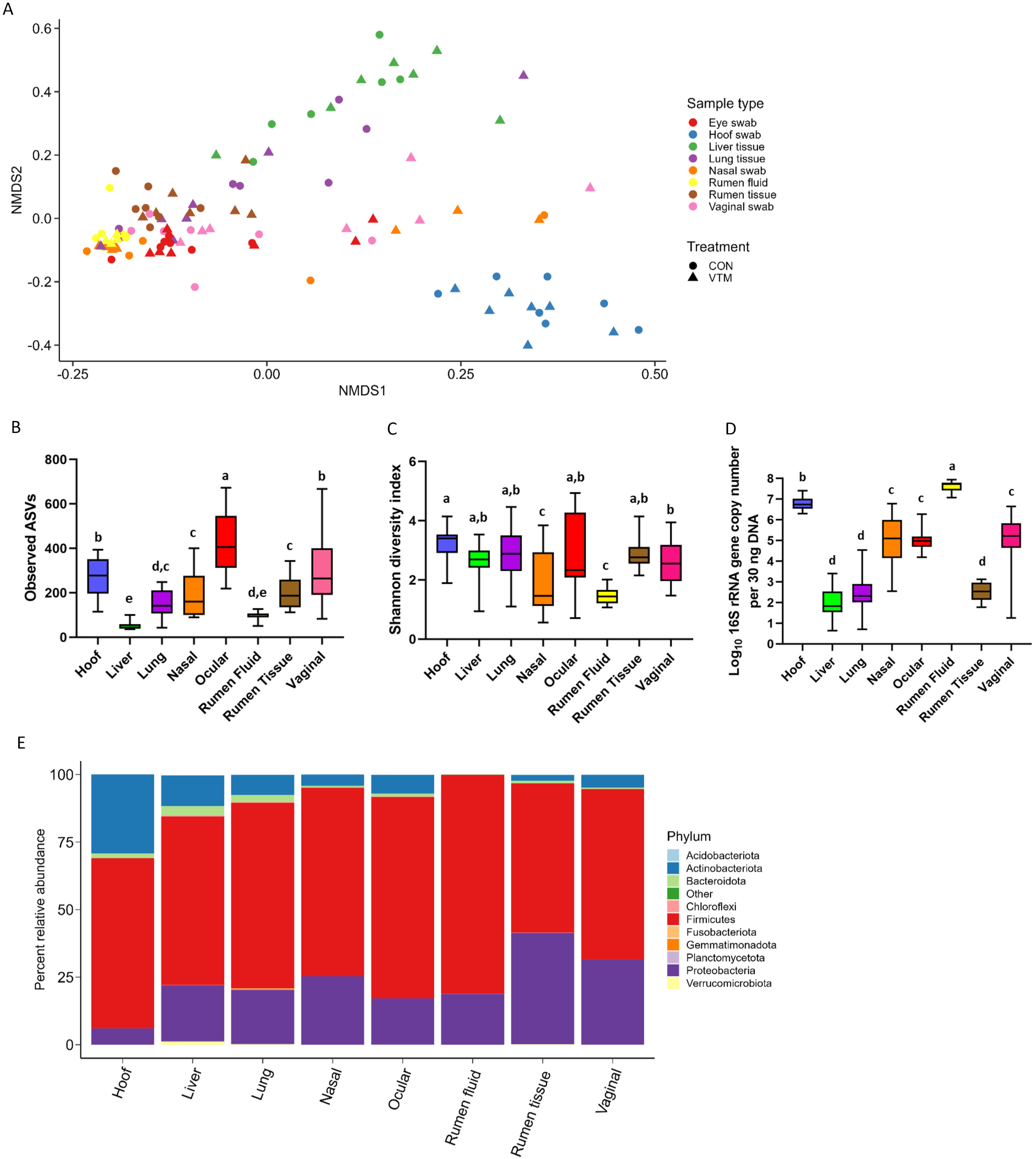
Comparison of the overall microbial community structure, richness, diversity, composition, and total microbial abundance of microbiota associated with 8 different newborn calf body sites (N = 14). (A) Non-metric multidimensional scaling (NMDS) plots of the Bray-Curtis dissimilarities; (B) Overall observed amplicon sequence variants (ASVs); (C) Overall Shannon diversity index; (D) Total bacterial abundance estimated by qPCR; (E) Relative abundance (%) of most relatively abundant bacterial phyla.

Fifteen genera that were differentially abundant between different sampling sites are presented in Fig. 9. The hoof microbiota harbored a greater relative abundance of *Corynebacterium, Macrococcus*, *Facklamia,* and *Peptostreptococcus* compared to all other sample types (*P* < 0.05). *Streptococcus* and *Clostridium sensu stricto* 7 were more relatively abundant in the ocular microbiota and the liver tissue-associated microbiota contained a greater relative abundance of *Staphylococcus* spp. The genus *Klebsiella* and *Escherichia Shigella* were more abundant in ruminal tissue, ruminal fluid, and vaginal microbiota.

**Figure 9.**
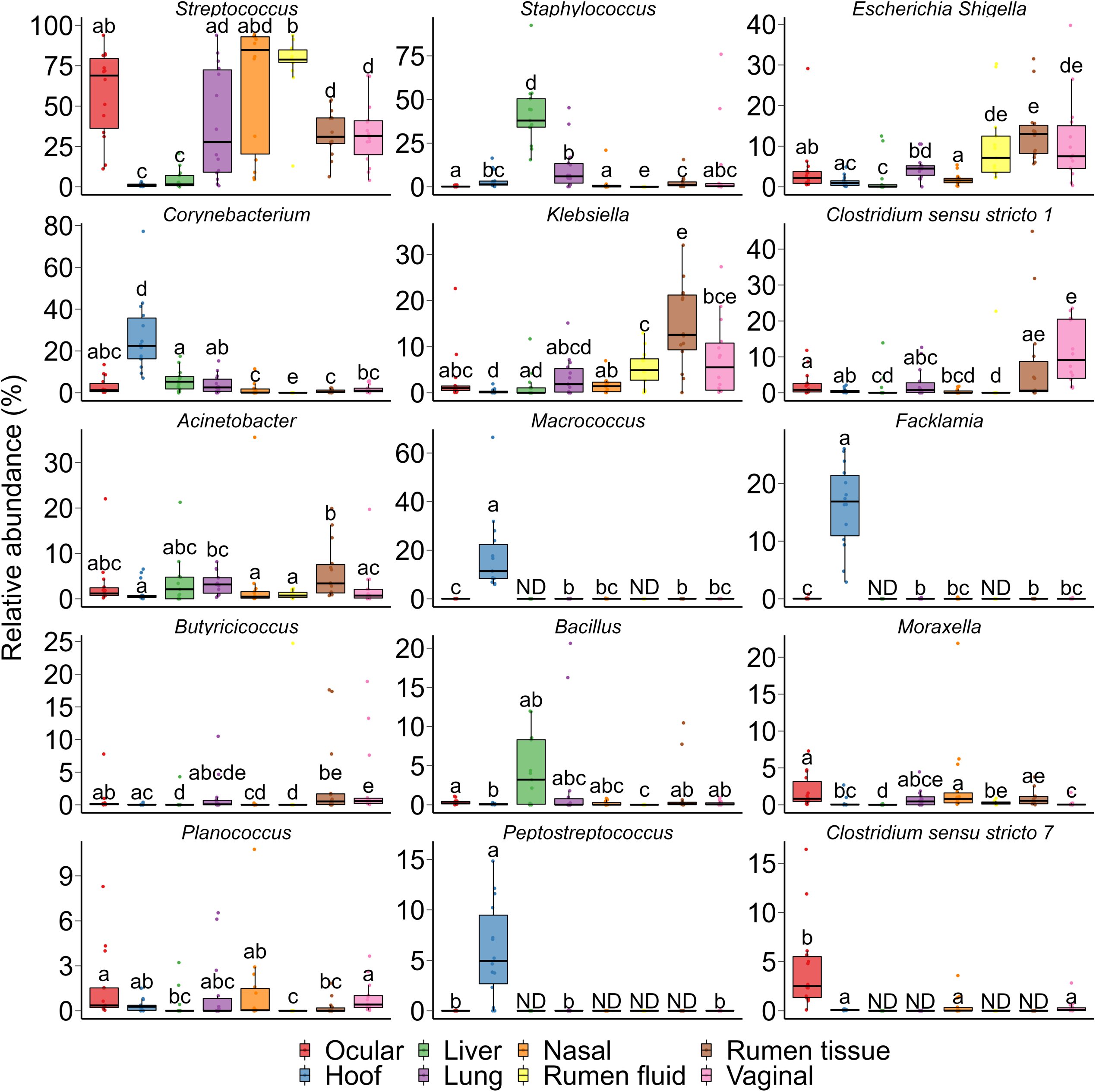
Bacterial genera that were differentially abundant between different sampling sites (N = 14 per sample type). ND denotes not detected; Different lowercase letters indicate significantly different means (*p* < 0.05).

According to the heatmap of the 100 most abundant ASVs (Supplementary Fig. S1), there was considerable inter-individual variation in both the prevalence and abundance of the major taxa present in the hoof, liver, lung, nasal, ocular, ruminal fluid, ruminal tissue, and vaginal samples. No single taxon was identified as being the most abundant between all 8 sampling locations. Several ASVs within the *Acinetobacter*, *Bibersteinia*, *Escherichia-Shigella*, *Klebsiella*, and *Streptococcus* genera were more abundantly present in nasal, ocular, ruminal fluid, ruminal tissue, and vaginal samples. Six ASVs within the *Aerococcaceae* family and *Facklamia* genus were exclusively present in ocular samples. Overall, most of these abundant ASVs were absent or only sparsely present in liver tissue.

### Effects of maternal vitamin and mineral supplement on immune response

Among the 15 tested serum cytokines (IFNγ, IL-1α, IL-1β, IL-4, IL-6, IL-8, IL-10, IL-17A, IL-36RA, IP-10, MCP-1, MIP-1α, MIP-1β, TNFα, and VEGF-A), concentrations of three cytokines were divergent in response to prenatal VTM supplementation (Fig. 10). The concentration of IP-10 was increased (*P* = 0.02) while IL-4 (*P* = 0.056) and IL-17A (*P* = 0.087) tended to be reduced in VTM calves compared to control calves.

**Figure 10.**
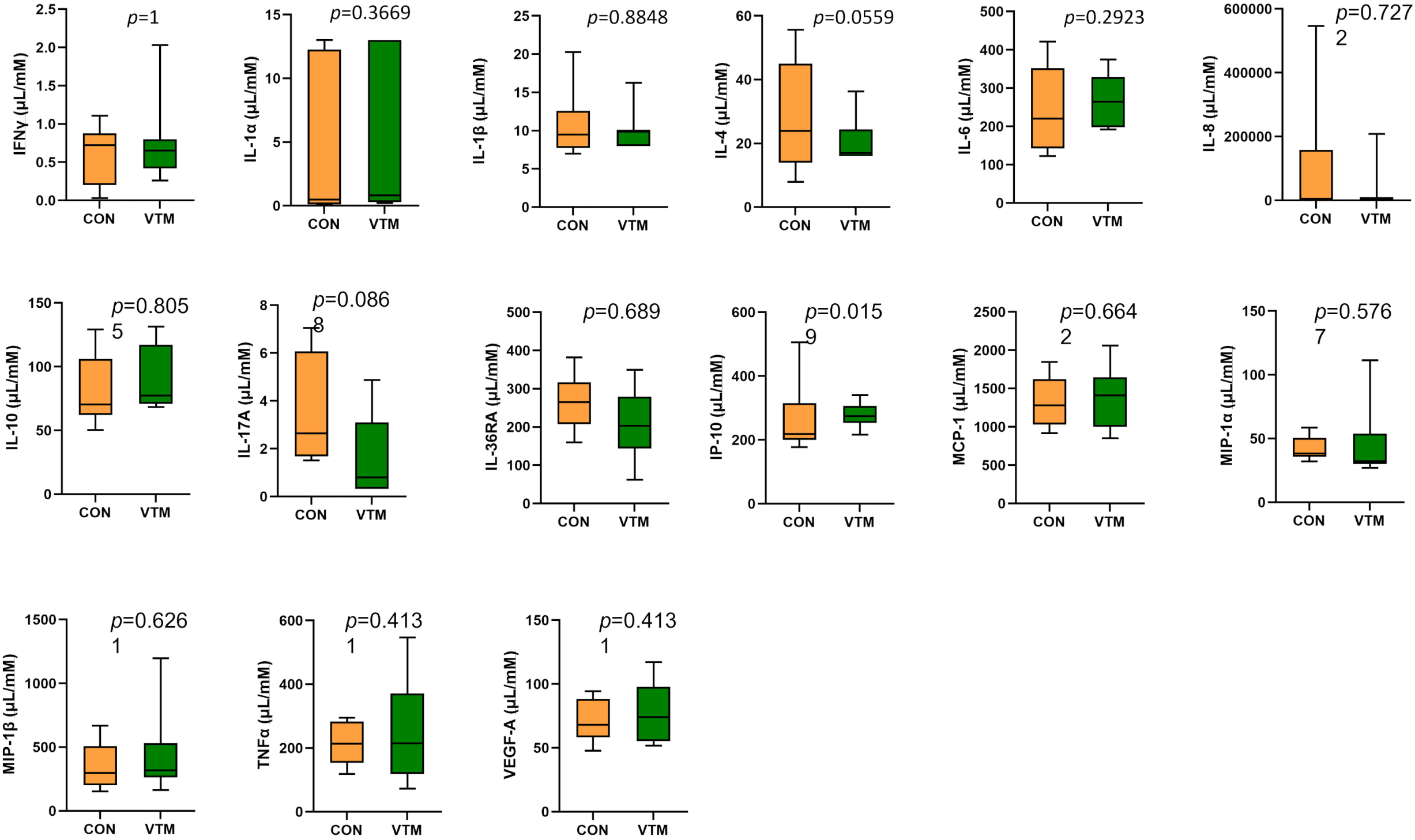
Observed cytokine concentration of newborn calves born from dams received either vitamin and mineral (VTM) supplementation or no VTM supplementation received (CON) sampled at 32 h after birth (n = 7/group).

## DISCUSSION

Although the neonatal gut harbors a low-diversity microbiota at birth, it progresses through various developmental, transitional, and stable phases before reaching an adult-like microbiota (53–55). Microbial colonization of the neonate sets the stage for the adult microbiome and the early-life microbiome is involved in the regulation of immune, endocrine, and metabolic developmental pathways (56), imparting long-term impacts on health and well-being later in life (57, 58). Recent evidence supporting *in utero* microbial colonization (19–21), and increased appreciation of the involvement of the microbiome in developmental programming (31, 59) highlights that maternal factors such as nutrition, act as modulators during pregnancy and could affect colonization and development of the fetal/neonatal/infant gut microbiota, thereby affecting the health and microbiome composition of offspring (60, 61).

While the impact of maternal nutrition on fetal programming and offspring development in cattle has received dedicated attention from researchers (27, 62, 63), the effect of maternal nutrition on the offspring microbiome remains largely unexplored. Therefore, in this study, we characterized 8 different microbial communities within and on calves to investigate whether the initial colonization of the microbiota in these anatomical locations were influenced by maternal VTM supplementation during pregnancy. The results of this study also provide a holistic view on the similarity and differences in the bacterial composition across the different body sites of newborn calves.

Average body weight of the dams at calving, and calf birth weight were not different (*P* > 0.05) between the VTM and non-VTM groups (Hurlbert et al., unpublished data). All calves at birth appeared to be normal and healthy. However, we observed that the ruminal, vaginal, and ocular microbiota were affected by maternal VTM supplementation. Following prenatal VTM supplementation, the offspring microbial community in the rumen (ruminal fluid and tissue-associated microbiota) diverged in terms of community structure (ruminal fluid), diversity (ruminal fluid and tissue), and composition at both the phylum and genus level. This suggests that maternal VTM supplementation during fetal development alters the initial microbial colonization of the neonatal calf gut. Since all calves were separated from the dam immediately at birth and were fed colostrum replacer (≤ 2 h) rather than being allowed to nurse, differences observed in the ruminal microbiota between the control and VTM calves occurred before and during birth. As such, we can rule out any influence on the neonate rumen microbiota from maternal milk, which is one of the microbial seeding sources for the neonatal gut (64, 65). One of the observed changes in the neonatal calf ruminal microbiota associated with prenatal VTM supplementation included increased microbial diversity. Lower diversity of the ruminal microbiota has previously been associated with greater feed efficiency (66, 67), and conversely, no link between the ruminal microbiota diversity and feed efficiency phenotype in weaned or older beef cattle has been reported (68, 69).

The ruminal microbiota in neonates undergoes dramatic changes in community structure, richness, and composition during the first 6 weeks of life, followed by more gradual alterations from 6 weeks to 3 months before development of an adult-like ruminal microbiota (70–73). It is difficult to predict the association of enhanced ruminal tissue microbiota diversity observed in VTM calves with future health and feed efficiency in adult life. However, the changes observed in microbial composition in the ruminal microbiota of VTM calves point out the positive association with calf gut health and feed efficiency. For example, VTM calves had an increased relative abundance of the phylum Bacteroidota, which has been reported to be positively associated with feed efficiency in cattle (69), as well as *Lactobacillus* spp. which have been reported to be more abundant in healthy calves (74) and have been shown to increase weight gain and improve feed conversion ratios while decreasing diarrhea in newborn calves when given as a probiotic (74–76). The *Escherichia-Shigella* genus contains potentially pathogenic species and strains and is positively associated with diarrhea in calves (74), and here it was reduced in VTM calves compared to control calves.

Although the community structure and diversity of the vaginal microbiota did not differ between the VTM and control calves, prenatal VTM supplementation increased microbial richness, altered the relative abundance of Bacteroidota, and reduced the total bacterial load in the vaginal mucosal surface. This suggests that VTM supplementation during pregnancy can influence the composition and community richness of the vaginal microbiota in neonatal calves. Compared to the gut microbiota, there is limited information available on the early life female reproductive microbiota in either humans or cattle, making it difficult to make any inferences on whether or how prenatal VTM supplementation-induced changes in the vaginal microbiota of calves would impact reproductive health and fertility later in life. Nevertheless, given the increased appreciation of the role of female reproductive microbiome in defining human (77, 78) and bovine reproductive health and fertility (5), our findings can be interpreted to imply that maternal nutritional factors could influence offspring reproductive tract microbiome.

A significant shift in terms of total bacterial abundance and community diversity were detected in the ocular microbiota of VTM calves. To the best of our knowledge, the microbial community associated with the eye is among the least characterized communities in cattle. However, the bovine ocular microbial community is a potential new target for manipulation, due to diseases such as IBK (79), which is a significant health issue affecting the beef cattle industry. Human studies suggest that the ocular surface microbiota is associated with ocular health (80, 81), and that a healthy ocular microbiota may provide resistance against colonization from infectious agents (82, 83). While the association of the ocular microbiota with IBK is yet to be determined, a recent study revealed a relative greater abundance of *Moraxella* spp. and *Mycoplasma* spp. in the bovine ocular microbiota, both of which are pathogens associated with IBK (7). This study also indicated that these opportunistic bacterial pathogens are common inhabitants of the ocular microbiota in beef calves ranging from 2-7 months old (7). This is supported by our data, as *Moraxella* was among the relatively abundant genera in the newborn calf ocular microbiota. Whether enrichment of *Moraxella* in the eye during early life can predispose calves to IBK or confer resistance against pathogenic *Moraxella* strains later in life is an important question that warrants further investigation. Microbial seeding of the calf oculus by dam’s vaginal microbiota during vaginal birth could be one of the sources accounting for the bacterial signature observed in oculus of these 32-h old calves. Overall, our results on the ruminal, vaginal, and ocular microbiota suggest that maternal VTM supplementation throughout pregnancy may have a niche-specific impact on neonatal microbial colonization. However, this is further verified by future studies.

Prenatal VTM supplementation-associated changes in maternal microbial and metabolic and immune profiles that induce alterations in the uterine microbial community may affect the colonization of the intestine, vagina, and ocular surface of calves born from VTM dams. In addition, vertical microbial transfer from the dam’s vagina to calves during birth (84–86) could not be ruled out as a potential factor influencing the newborn microbiota. However, the extent of vertical microbial transfer during birth and its contributions to the distinctive compositional and diversity changes in calves born from VTM supplemented dams remain to be explored.

Although no treatment effect was detected in the nasal, lung, liver, and hoof microbiota, our data on the composition of respiratory, hepatic, and hoof microbiota are novel, given that the newborn calves had only minimal exposure to microbes from the dam (no milk or colostrum, and minimal physical contact). Interestingly, we identified several bacterial genera that include pathogens such as *Mannheimia* (*Mannheimia haemolytica;* BRD) (87), *Moraxella* (BRD and IBK), and *Fusobacterium* (*Fusobacterium necrophorum*; bovine liver abscesses and bovine digital dermatitis) (88, 89) in the hoof, nasal, and lung microbiota of our healthy newborn calves. This poses an important question as to whether the initial colonization with these pathogenic bacteria in the respiratory tract and on hooves could have adverse or beneficial impacts on the offspring microbiome and immune development, and their resistance to BRD, liver abscesses, and hoof infections later in life. These bacterial genera, however, were not relatively abundant in the liver tissue of the newborn calves. The liver has traditionally been believed to be sterile;however, this has recently been challenged by the identification of a commensal microbiota in the liver of healthy cattle (90) and humans (91, 92).

As expected, the overall microbial structure, diversity, composition, and bacterial abundance were noticeably different among the microbiota characterized, despite being dominated by only five bacterial phyla. Several factors such as niche-specific physiological factors (temperature, pH, oxygen, and nutrient availability), diet, and environment are involved in shaping the respiratory (93, 94), gut (4) and reproductive tract (95) microbiota. Subtle physiological and anatomical differences in the mucosal surfaces of the bovine respiratory tract can shift the bacterial composition along the bovine respiratory (96) and GI tract (86). However, the presence of members from certain beneficial genera, such as *Lactobacillus,* or potentially pathogenic genera including *Moraxella* and *Mannheimia*, in almost all anatomical sites highlight that there may be transfer of microbes between different body sites. The same bacterial species present in multiple body sites in calves/cattle may indicate site-specific functions and adaptation. Such body site-specific genome adaptations have been reported in *Lactobacillus* spp. (97). Shotgun metagenomic sequencing is necessary to evaluate the whole-body microbiome of cattle to identify pathogenic bacterial species and strains that are shared amongst these different body sites and their site-specific functions. This would provide a holistic view of managing bovine pathogenic bacteria in a more effective and targeted manner.

Nutritional supplementation including VTM supplementation during pregnancy (98) and maternal gut microbiome alterations via nutrient supplementation or antibiotic usage (65, 99) can have an impact on neonatal immune system development. The comparison of serum concentrations of 15 different cytokines between the two group calves in the present study showed that maternal VTM supplementation during gestation may have minimal impact on neonatal serum cytokine profiles. Serum IP-10, IL-4 and IL-17A in neonatal calves responded to prenatal VTM supplementation. Interferon gamma-induced protein 10 (IP-10), which is secreted from T lymphocytes, neutrophils, monocytes, endothelial cells, and fibroblasts (100), is involved in inflammatory responses in various infectious diseases (101, 102). IP-10 was elevated in the serum of VTM calves. Conversely, VTM calves tended to have lower serum concentrations of proinflammatory cytokines IL-4 and IL-17A, both of which have significant immune regulator functions that are critical in mediating inflammatory and infectious responses (103–105). The other 13 evaluated cytokines (IFNγ, IL-1α, IL-1β, IL-6, IL-8, IL-10, IL-36RA, MCP-1, MIP-1α, MIP-1β, TNFα, and VEGF-A) did not respond to maternal VTM supplementation. This might be due to smaller sample size, or it is possible that these specific cytokines are not sensitive to changes in vitamin and mineral status. Nevertheless, our results indicate that VTM supplementation during fetal development may have minimal but specific effects on the fetal and neonatal calf immune system. The impact of maternal VTM supplementation on the ruminal, vaginal, and ocular microbiota of neonatal calves begs a question to whether prenatal VTM supplementation could influence localized immune system development in the gut, reproductive tract, and the eye in offspring. In addition, evaluation of other immune indicators, such as passive transfer of immunity and titer response to vaccination at later post-natal time points is warranted.

## CONCLUSION

Although maternal VTM supplementation during pregnancy had minimal impact on the nasal, lung, liver, and hoof microbiota of the neonatal calf, the ruminal, vaginal, and ocular microbiota appeared to be affected. The respiratory (nasal and lung), ocular, vaginal, and ruminal fluid microbiota were most similar while the hoof and liver microbiota were most dissimilar to the other microbiota. Some genera, such as those that include pathogens associated with BRD, liver abscesses, IBK, and digital dermatitis were present in almost all anatomical sites evaluated. Overall, our study indicates that the gut, hoof, ocular, liver, respiratory, and reproductive sites of newborn calves are colonized by a diverse, dynamic, and site-specific microbiota, and that prenatal VTM supplementation may influence the gut, vaginal and ocular microbiota of newborns, and have minimal influence on blood cytokine profiles. These findings are important for directing future research on developing maternal nutrition and microbiome targeted approaches to improve offspring microbiome development, feed efficiency and disease resilience.

## DATA AVAILABILITY

Raw sequence data are available from the NCBI Sequence Read Archive under BioProject accession PRJNA890

## ACKNOWLEDGEMENTS

Microbiological work presented in this study was financially supported by the North Dakota Agricultural Experiment Station as part of a start-up package for S.A. Animal procurement and management portion of the project was supported by the North Dakota State Board of Agricultural Research and Education (SBARE), the Central Grasslands Research and Extension Center, and the ND Agriculture Experiment Station. We acknowledge the support from the employees of the NDSU Beef Cattle Research Complex and Animal Nutrition and Physiology Center.

## AUTHOR CONTRIBUTIONS

Conceiving the idea, designing the study, and providing supervision—S.A. and C.R.D.; cattle management: C.R.D., K.K.S., J.L.H., F.B, and A.C.B.M.; Animal care and sample collections--S.A., C.R.D., K.C.S., J.L.H., A.C.B.M., K.A.B., J.D.K. and F.B.; Sample processing---K.N.S., S.M.L.; Data processing, bioinformatics and statistical analysis---D.B.H., K.J., S.A. S.M.L; Manuscript writing: S.M.L and S.A.; Manuscript review, editing and finalizing---S.M.L., S.A., D.B.H., C.R.D., and K.C.S; All authors have read and agreed to the published version of the manuscript.

## COMPETING INTERESTS

The authors declare no competing interest.

