## Supplementary Tables 1, 3, 4, 5 and Fig. S1 for "Whole-body Microbiota of Newborn Calves and Their Response to Prenatal Vitamin and Mineral Supplementation"

**Supplementary Table S1.** Mineral and vitamin supplement composition.

| **Nutrient** | **Assurance levels** | |
|---|---|---|
| ***Minerals*[1]** | **Min** | **Max** |
| Calcium, g / kg of DM | 135.0 | 162.0 |
| Phosphorus, g / kg of DM | 75.0 | - |
| Sodium cholride, g / kg of DM | 180.0 | 216.0 |
| Magnesium, g / kg of DM | 10.0 | - |
| Potassium, g /kg of DM | 10.0 | - |
| Manganese, mg / kg of DM | 3,600.0 | - |
| Cobalt, mg / kg of DM | 12.0 | - |
| Copper, mg / kg of DM | 1200.0 | - |
| Iodine, mg / kg of DM | 60.0 | - |
| Selenium, mg / kg of DM | 27.0 | - |
| Zinc, mg / kg of DM | 3,600.0 | - |
| ***Vitamins, IU / kg of DM*** | | |
| A | 661,500.0 | |
| D | 66,150.0 | |
| E | 661.5 | |

[1]Purina Wind and Rain Storm All Season 7.5 Complete Mineral (Land O' Lakes, Inc., Arden Hills, MN); ingredients: dicalcium phosphate, monocalcium phosphate, processed grain by-products, plant protein products, calcium carbonate, molasses products, salt, mineral oil, potassium chloride, magnesium oxide, ferric oxide, vitamin E supplement, vitamin A supplement, lignin sulfonate, cobalt carbonate, manganese sulfate, ethylenediamine dihydroiodide, zinc sulfate, copper chloride, vitamin D3 supplement, natural and artificial flavors, and sodium selenite. DM = drive matter.

**Supplementary Table S3**. Average number of sequencing reads (±SEM) before and after removing removal of mitochondria, chloroplast, and negative control sequences.

| **Sample type** | **Sequencing reads before removal of potential contaminant sequencing reads** | **Sequencing reads after removal of potential contaminant sequencing reads** |
|---|---|---|
| Hoof swab | 185904 ± 9970 | 185261 ± 10017 |
| Liver tissue | 232797 ± 18164 | 89852 ± 10847 |
| Lung tissue | 484801 ± 38142 | 369539 ± 42475 |
| Nasal swab | 257610 ± 31260 | 254352 ± 32171 |
| Ocular swab | 272472 ± 20448 | 264921 ± 20423 |
| Ruminal fluid | 276086 ± 16693 | 275955 ± 16704 |
| Ruminal tissue | 432122 ± 26428 | 400929 ± 30425 |
| Vaginal swab | 225180 ± 14767 | 223692 ± 15009 |

**Supplementary Table S4**. PERMANOVA analysis summary for comparison of microbial community structure between different sampling type.

| PERMANOVA | $R^2$ | Adjusted *P - value* | PERMANOVA | $R^2$ | Adjusted *P - value* |
|---|---|---|---|---|---|
| Ruminal fluid vs. Hoof swab | 0.64 | 0.0028 | Liver tissue vs. Vaginal swab | 0.301 | 0.0028 |
| Ruminal fluid vs. Liver tissue | 0.54 | 0.0028 | Liver tissue vs. Ocular swab | 0.434 | 0.0028 |
| Ruminal fluid vs. Ruminal tissue | 0.33 | 0.0028 | Ruminal tissue vs. Vaginal swab | 0.121 | 0.0028 |
| Ruminal fluid vs. Vaginal swab | 0.25 | 0.0028 | Ruminal tissue vs. Ocular swab | 0.257 | 0.0028 |
| Nasal swab vs. Hoof swab | 0.41 | 0.0028 | Ruminal fluid vs. Lung tissue | 0.181 | 0.0448 |
| Nasal swab vs. Liver tissue | 0.35 | 0.0028 | Ruminal fluid vs. Ocular swab | 0.172 | 0.0168 |
| Nasal swab vs. Ruminal tissue | 0.23 | 0.0028 | Nasal swab vs. Vaginal swab | 0.142 | 0.0448 |
| Hoof swab vs. Lung tissue | 0.39 | 0.0028 | Lung tissue vs. Ruminal tissue | 0.140 | 0.0168 |
| Hoof swab vs. Liver tissue | 0.45 | 0.0028 | Vaginal swab vs. Ocular swab | 0.147 | 0.0196 |
| Hoof swab vs. Ruminal tissue | 0.52 | 0.0028 | Ruminal fluid vs. Nasal swab | 0.127 | 0.493 |
| Hoof swab vs. Vaginal swab | 0.38 | 0.0028 | Nasal swab vs. Lung tissue | 0.096 | 0.580 |
| Hoof swab vs. Ocular swab | 0.51 | 0.0028 | Nasal swab vs. Ocular swab | 0.053 | 1.00 |
| Lung tissue vs. Liver tissue | 0.20 | 0.0028 | Lung tissue vs. Vaginal swab | 0.099 | 0.11 |
| Liver tissue vs. Ruminal tissue | 0.40 | 0.0028 | Lung tissue vs. Ocular swab | 0.123 | 0.11 |

**Supplementary Table S5**. Percent relative abundance (%) of list of these bacteria genera that have been reported to encompasses species associated with bovine infectious diseases*.

| Genus | Rank | CON | VTM | SEM |
|---|---|---|---|---|
| **Ocular swab** | | | | |
| *Moraxella* | 8 | 0.73 | 3.07 | 0.62 |
| *Mannheimia* | 20 | 0.05 | 0.6 | 0.17 |
| *Fusobacterium* | 255 | 0.005 | 0.003 | 0.004 |
| *Trueperella* | 301 | 0 | 0.006 | 0.003 |
| *Histophilus* | 372 | 0 | 0.003 | 0.001 |
| **Liver tissue** | | | | |
| *Pandoraea* | 16 | 0.78 | 0.84 | 0.57 |
| *Haemophilus* | 18 | 0.23 | 1.15 | 0.46 |
| *Megasphaera* | 35 | 0.86 | 0 | 0.43 |
| *Ochrobactrum* | 38 | 0.81 | 0 | 0.4 |
| *Thermicanus* | 39 | 0 | 0.68 | 0.24 |
| *Pelomonas* | 40 | 0.32 | 0.39 | 0.3 |
| **Nasal swab** | | | | |
| *Moraxella* | 4 | 1.45 | 4.35 | 1.91 |
| *Mannheimia* | 10 | 1.82 | 1.16 | 1.22 |
| *Fusobacterium* | 121 | 0.017 | 0 | 0.008 |
| **Vaginal swab** | | | | |
| *Mannheimia* | 12 | 1.1 | 0.96 | 0.8 |
| *Moraxella* | 27 | 0.02 | 0.32 | 0.12 |
| *Fusobacterium* | 361 | 0.0004 | 0.0007 | 0.0004 |
| *Trueperella* | 447 | 0.0004 | 0 | 0.0002 |
| **Hoof swab** | | | | |
| *Mannheimia* | 13 | 1.84 | 0.16 | 0.98 |
| *Moraxella* | 26 | 0.42 | 0.17 | 0.26 |
| *Fusobacterium* | 37 | 0.2 | 0.01 | 0.08 |
| *Trueperalla* | 79 | 0.03 | 0.005 | 0.015 |
| **Lung tissue** | | | | |
| *Moraxella* | 13 | 0.6 | 1.09 | 0.44 |
| *Mannheimia* | 28 | 0.01 | 0.63 | 0.29 |
| *Fusobacterium* | 40 | 0.4 | 0.01 | 0.2 |
| *Trueperella* | 228 | 0.009 | 0 | 0.005 |
| **Rumen fluid** | | | | |
| *Moraxella* | 9 | 0.34 | 0.28 | 0.09 |
| *Mannheimia* | 12 | 0.03 | 0.13 | 0.06 |
| **Rumen tissue** | | | | |
| *Moraxella* | 11 | 0.95 | 1.17 | 0.52 |
| *Mannheimia* | 25 | 0.07 | 0.28 | 0.13 |
| *Fusobacterium* | 163 | 0.015 | 0 | 0.0075 |

*CON: calves born from non-VTM supplemented dams (n = 7); VTM: calves born from VTM supplemented dams (n =7).  SEM: standard error of the mean.

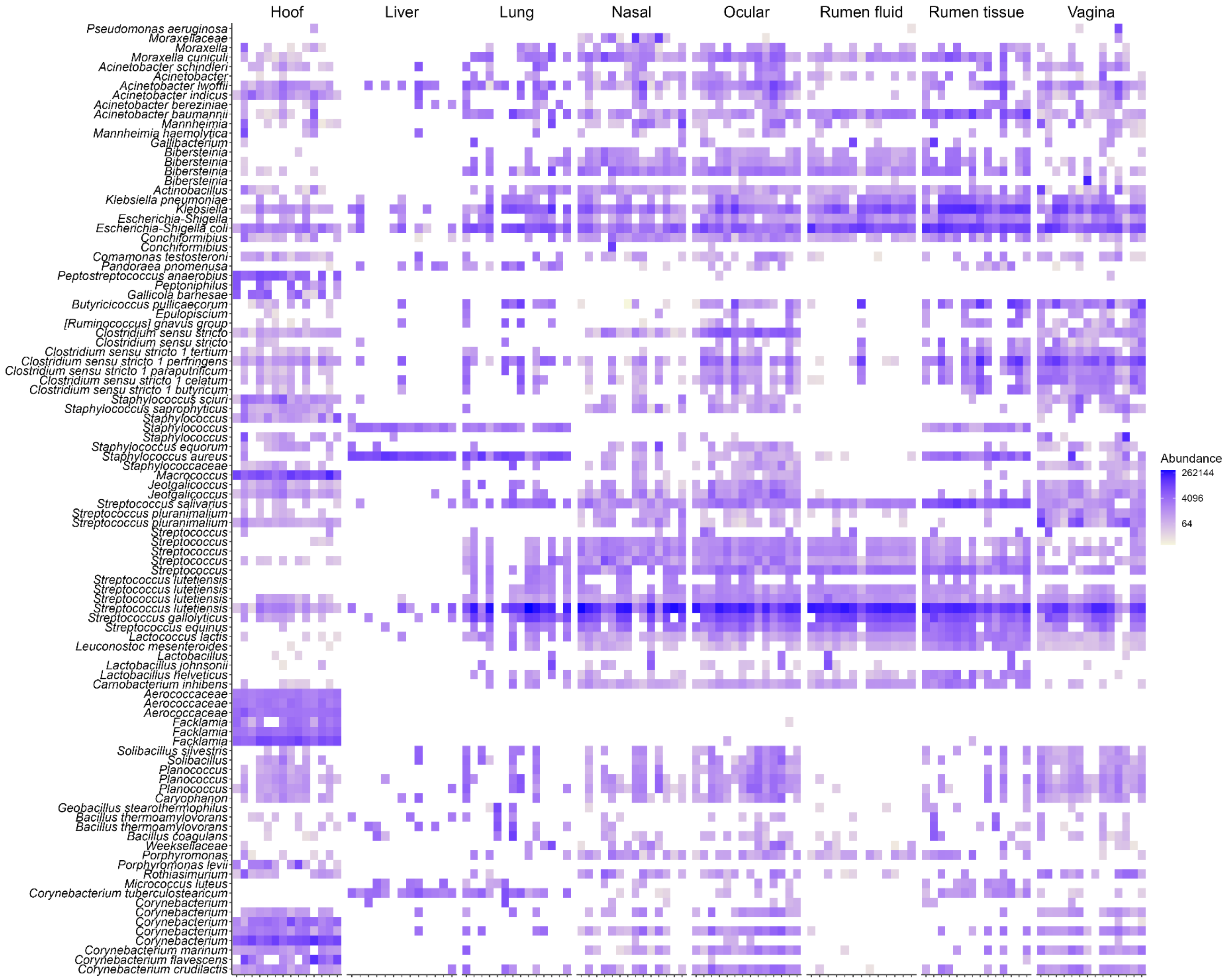


**Supplementary Figure S1**. Heatmap showing the 100 most abundant ASVs (log4) overall sample type.
